# *vcf2gwas* - Python API for comprehensive GWAS analysis using GEMMA

**DOI:** 10.1101/2021.06.01.446586

**Authors:** Frank Vogt, Gautam Shirsekar, Detlef Weigel

## Abstract

We present a new software package *vcf2gwas* to perform reproducible genome-wide association studies (GWAS). *vcf2gwas* is a Python API for bcftools, PLINK and GEMMA. Before running the analysis a traditional GWAS workflow requires the user to edit and format the genotype information from commonly used Variant Call Format (VCF) file and phenotype information. Post-processing steps involve summarizing and visualizing the analysis results. This workflow requires a user to utilize the command-line, manual text-editing and knowledge of one or more programming/scripting languages which can be time-consuming especially when analyzing multiple phenotypes. Our package provides a convenient pipeline performing all of these steps, reducing the GWAS workflow to a single command-line input without the need to edit or format the VCF file beforehand or to install any additional software. In addition, features like reducing the dimensionality of the phenotypic space and performing analyses on the reduced dimensions or comparing the significant variants from the results to specific genes/regions of interest are implemented. By integrating different tools to perform GWAS under one workflow, the package ensures reproducible GWAS while reducing the user efforts significantly.

**Availability and implementation:** The source code of *vcf2gwas* is available under the GNU General Public License. The package can be easily installed using conda. Installation instructions and a manual including tutorials can be accessed on the package website at https://github.com/frankvogt/vcf2gwas

## 1 Introduction

Genome-wide association study (GWAS) has been proven to be an extremely useful tool to find an association between genetic variants, typically single-nucleotide polymorphisms (SNPs) and a given trait in many organisms. To carry out a GWAS, the traits under investigation are analyzed by recording the phenotypes of different individuals and sequencing their DNA. After the SNPs are read from the genotype information, these genetic variants are tested for the association with the phenotypes using various computational methods implemented in a wide range of software. The likelihood of each SNP to be associated with the trait is calculated and subsequently used to identify SNPs with significant associations.

While there are many algorithms implemented in various software to perform the association analysis, Genome-wide Efficient Mixed Model Association (GEMMA) ([1]) stands out because of its versatility and efficiency in handling large scale data. GEMMA can fit a univariate linear mixed model ([1]), a multivariate mixed model ([2]), and a Bayesian sparse linear mixed model ([3]) for testing marker associations with a trait of interest in different organisms.

Although GEMMA has a very straightforward command-line interface to carry out the actual association analysis, it requires users to first install necessary software with required dependencies on their machines. Subsequently, users need to prepare the inputs (genotype and phenotype) in a proper format before execution of GEMMA. Similarly, the outputs generated need post-processing of the results for better interpretation and presentation. The entire workflow can be overwhelming especially for inexperienced users. Briefly, this workflow starts with the genotype information in a VCF file. This file has to be converted to the PLINK[4] BED format with the phenotype information that needs to be manually edited into associated .FAM file. Once the analysis is complete, the user is left with the outputs which need to be summarized and plotted. If multiple phenotypes are to be analyzed, the analysis has to be repeated for every phenotype. Thus, performing GWAS in this way using GEMMA is very time-consuming and makes it very challenging to conduct reproducible research while keeping all the analyses well-organized.

The *vcf2gwas* package aims to facilitate performing a GWAS with GEMMA by automating all the phases, beginning with the installation of all the required software, to input preparations, to carrying out the analysis, and finally to processing the results. This reduces overhead for the user to a minimum. *vcf2gwas* is especially helpful when analyzing large numbers of phenotypes or different sets of individuals because it can perform the analyses in parallel with a single .csv file with all the phenotypes. Additionally, the package offers features like analyzing reduced phenotypic space and comparing the SNPs showing significant association to genes or regions of interest. *vcf2gwas* is easily installable as a conda package and therefore makes it possible to reproduce each GWAS on any compatible machine.

## 2 Implementation

The *vcf2gwas* package is implemented using the Python programming language. It contains multiple Python3 scripts with all the functions required for the program to execute the analysis. The core functionality of the package consists of the ability to perform automated GWAS analysis with a single command-line input, requiring minimal overhead by the user. It has multiple wrapper functions calling bcftools ([5]), PLINK ([4]), and GEMMA ([1]) from Python3 as well as complementary Python3 functions performing both pre- and post-analysis tasks, all executed by the pipeline scripts. Pre-analysis includes functions for formatting, trimming and filtering the input files for the association analysis with GEMMA. The different analysis modes of GEMMA can be utilized and are also specified with the command-line input. For the post-analysis, additional functions are used to summarize and visualize the resulting data and perform additional operations specified by the command-line options. A tree-like directory structure is generated to save all the results associated with the analysis of a given phenotype. *vcf2gwas* uses Python libraries numpy[6], matplotlib[7], pandas[8] for preparing input/output files and plotting while scikit-learn[9] and umap-learn libraries are used for dimensionality reduction.

## 3 Example usage

Performing a GWAS using GEMMA requires a user to execute a number of steps. First, the genotype information (usually available as VCF file) has to be converted to the PLINK BED format. Second, phenotype information must be provided in a specific format as well, for e.g. by manually editing FAM files. Furthermore, the user has to trim both the genotype and phenotype information to contain the matching individuals in correct order. And finally, additional analyses such as visualizing and summarizing the results require the user to utilize additional software or programming languages. In addition, analyzing more than one phenotype results in re-running GEMMA for each phenotype.

In contrast, by running *vcf2gwas*, the pipeline will perform all of the above steps in a fully automated fashion. The input files will be trimmed, filtered and converted to the format required by GEMMA. After the analysis is executed, the results will be summarized and visualized by drawing publication-ready Manhattan (Figure 1) and Q-Q plots; optionally additional operations as explained in the next section can be carried out. A basic bash command example to analyze a single phenotype using GEMMA’s linear mixed model providing only the VCF and phenotype file is:

~~~
$ vcf2gwas -v <input.vcf> -pf <inputpheno.csv> -p 1 -lmm
~~~

**Figure 1:**
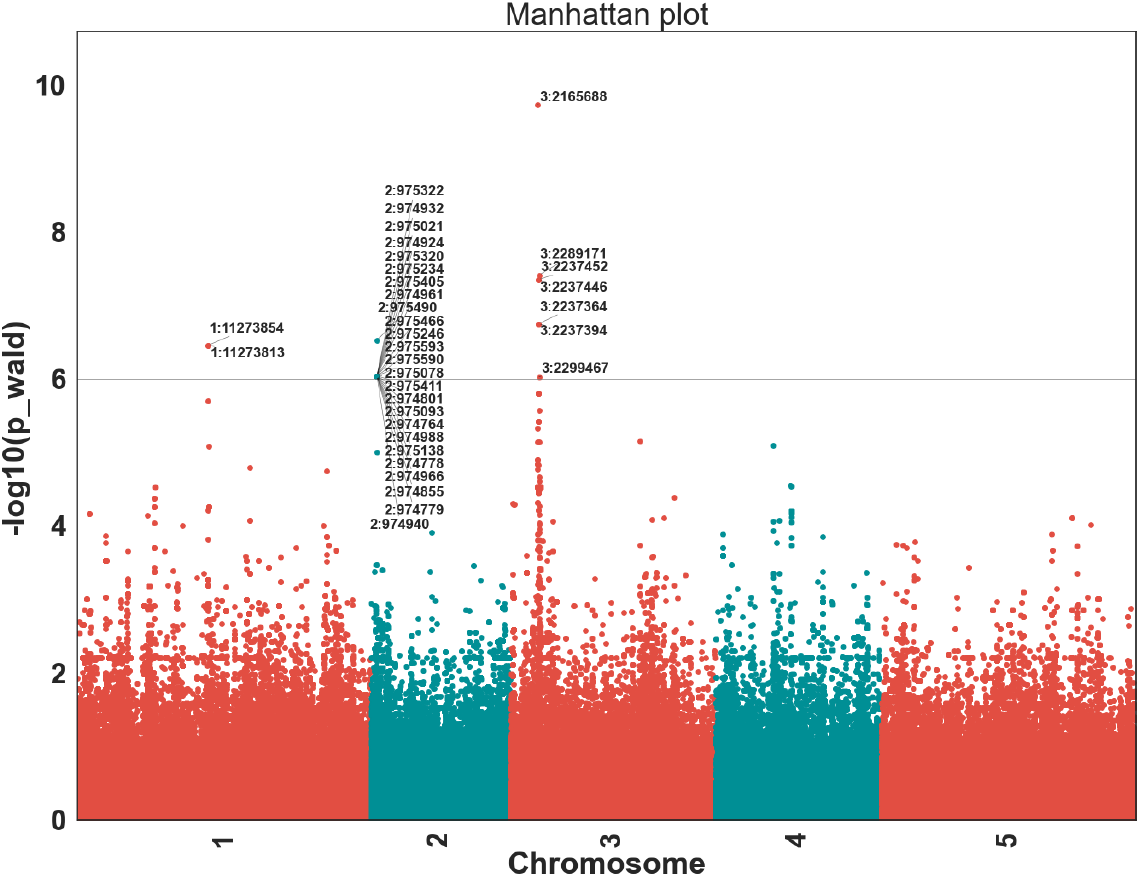
Manhattan plot produced by *vcf2gwas* on *avrRpm1* recognition in *Arabidopsis thaliana*. Linear mixed model analysis on a hypersensitive response phenotype observed in 58 *A. thaliana* host lines in response to *Pseudomonas syringe* expressing *avrRpm1* gene. The most significant SNP is 700 bp upstream of the *A. thaliana Rpm1* resistance gene. Description of the original experiment can be found at https://arapheno.1001genomes.org/phenotype/17/.

Internally, the pipeline converts and formats the input files utilizing bcftools and PLINK while the post-analysis steps are performed using custom Python functions. We have presented an example Manhattan plot of a GWA analysis performed with *vcf2gwas* in figure 1. Q-Qplot for the analysis is in Fig S4 and the list of the SNPs found nearby a given set of *Arabidopsis thaliana* NBS-LRR domain containing immune system genes with their coordinates is presented in Table S1.

## 4 Additional functionality

Beyond the basic necessities of summarizing and plotting the results of the GEMMA analysis, it might be desirable to perform the GWAS in reduced phenotypic space to explore genetic associations for underlying phenotypic structure or to compare the resulting relevant SNPs to genes/regions of interest. Below are the most important additional features implemented in *vcf2gwas*:

- Dimensionality reduction via PCA or UMAP can be performed on phenotypes and used for analysis.
- Relevant SNPs can be compared to genes/regions of interest listed in additionally specified file.
- *vcf2gwas* is able to analyze several input files with different sets of individuals and multiple phenotypes in a efficient manner due to parallelization, saving the user a lot of time compared to standard GWAS procedure.
- Results are reproducible on any Unix-based machine (macOS / Linux).

## Supplementary material

Supplementary data is available online.

## Acknowledgements

We thank Hajk-Georg Drost and Max Collenberg for useful comments and suggestions during the software development.

## Funding

This work has been supported by the Max Planck Society.

## Notes

### Competing Interest Statement

The authors have declared no competing interest.

