## Supplementary Data for "*vcf2gwas* - Python API for comprehensive GWAS analysis using GEMMA"

### Supplementary Figures

### Supplementary Tables

#### List of Supplementary Tables

1 List of SNPs above the threshold ($-log_{10}(10^{-7})$) and their distance from a given gene set (here, *Arabidopsis thaliana* immune system genes containing NBS-LRR domain) . 4

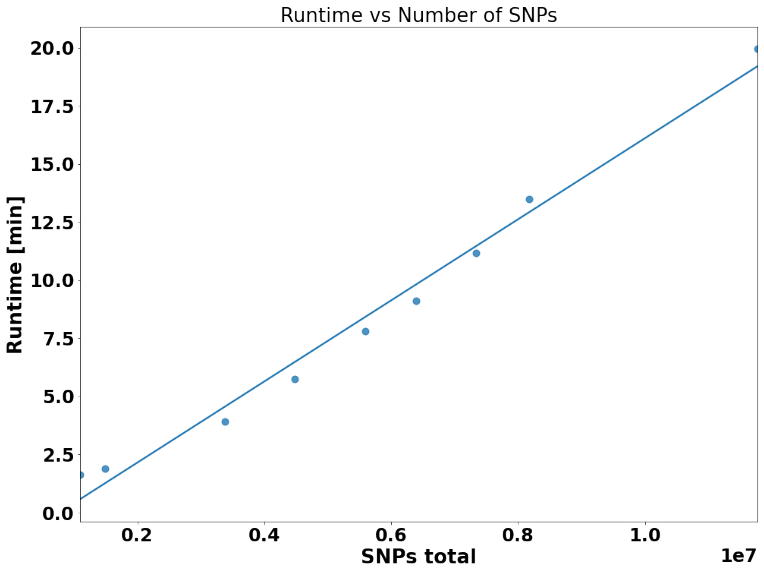

**Figure S 1:** *vcf2gwas* runtime as a function of different level of linkage disequilibrium (LD)-based pruning of the original SNP dataset.

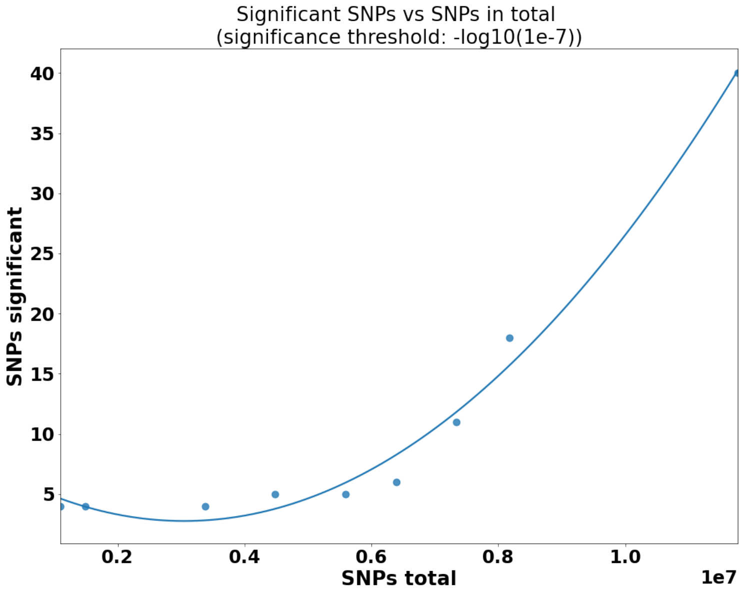

**Figure S 2:** Number of significant SNPs as a function of SNPs retained by different LD-pruning thresholds

The threshold to distinguish significant SNPs is $-log_{10}(10^{-7})$.

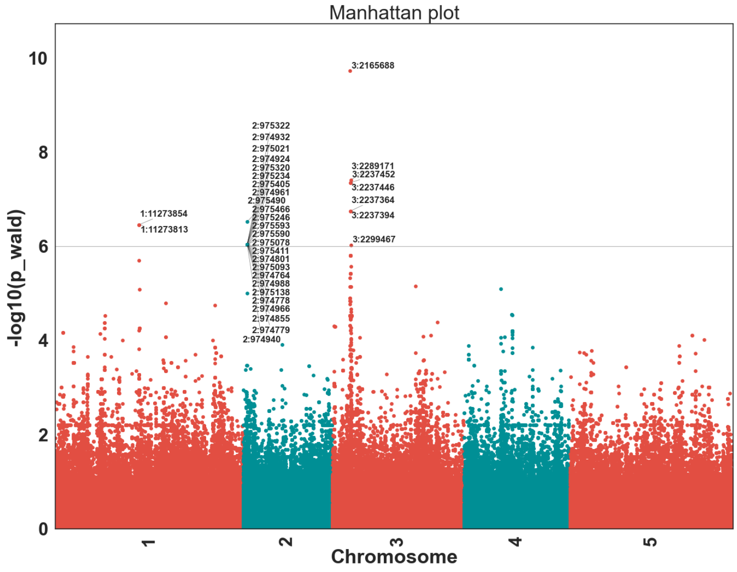

**Figure S 3:** Manhattan plot produced by *vcf2gwas* on association analysis performed on the *avrRpm1* recognition in *Arabidopsis thaliana*

Linear mixed model analysis on a hypersensitive response phenotype observed in 58 *A. thaliana* host lines in response to *Pseudomonas syringe* expressing *avrRpm1* gene. The most significant SNP is 700 bp upstream of the *A. thaliana Rpm1* resistance gene. Description of the original experiment can be found at `https://arapheno.1001genomes.org/phenotype/17/`.

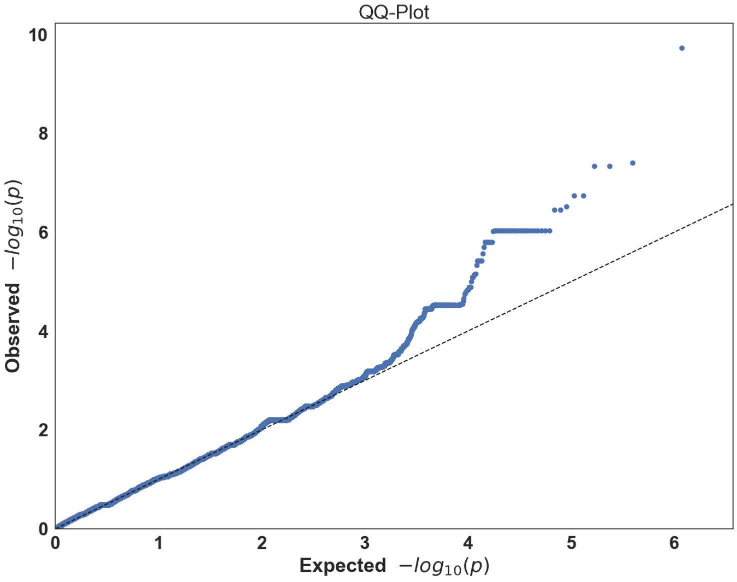

**Figure S 4:** Q-Q plot produced by *vcf2gwas* on association analysis performed on the *avrRpm1* recognition in *Arabidopsis thaliana*

| SNP_ID | chr | phenotypes | gene_ID(up) | gene_comment(up) | gene_name(up) | gene_distance(up) | SNP_pos | gene_distance(down) | gene_name(down) | gene_comment(down) | gene_ID(down) |
|---|---|---|---|---|---|---|---|---|---|---|---|
| 3:2237364 | 3 | avrRpm | AT3G07040.1 | NB-ARC domain-containing disease resistance protein | RPM1 | 8340 | 2237364 | | | | |
| 3:2237394 | 3 | avrRpm | AT3G07040.1 | NB-ARC domain-containing disease resistance protein | RPM1 | 8370 | 2237394 | | | | |
| 3:2237446 | 3 | avrRpm | AT3G07040.1 | NB-ARC domain-containing disease resistance protein | RPM1 | 8422 | 2237446 | | | | |
| 3:2237452 | 3 | avrRpm | AT3G07040.1 | NB-ARC domain-containing disease resistance protein | RPM1 | 8428 | 2237452 | | | | |
| 3:2289171 | 3 | avrRpm | AT3G07040.1 | NB-ARC domain-containing disease resistance protein | RPM1 | 60147 | 2289171 | | | | |
| 2:975138 | 2 | avrRpm | AT2G03030.1 | Toll-Interleukin-Resistance (TIR) domain family protein | | 84618 | 975138 | 28330 | | Toll-Interleukin-Resistance (TIR) domain family protein | AT2G03300.1 |
| 2:975234 | 2 | avrRpm | AT2G03030.1 | Toll-Interleukin-Resistance (TIR) domain family protein | | 84714 | 975234 | 28234 | | Toll-Interleukin-Resistance (TIR) domain family protein | AT2G03300.1 |
| 2:975320 | 2 | avrRpm | AT2G03030.1 | Toll-Interleukin-Resistance (TIR) domain family protein | | 84800 | 975320 | 28148 | | Toll-Interleukin-Resistance (TIR) domain family protein | AT2G03300.1 |
| 2:975405 | 2 | avrRpm | AT2G03030.1 | Toll-Interleukin-Resistance (TIR) domain family protein | | 84885 | 975405 | 28063 | | Toll-Interleukin-Resistance (TIR) domain family protein | AT2G03300.1 |
| 2:975411 | 2 | avrRpm | AT2G03030.1 | Toll-Interleukin-Resistance (TIR) domain family protein | | 84891 | 975411 | 28057 | | Toll-Interleukin-Resistance (TIR) domain family protein | AT2G03300.1 |
| 2:975490 | 2 | avrRpm | AT2G03030.1 | Toll-Interleukin-Resistance (TIR) domain family protein | | 84970 | 975490 | 27978 | | Toll-Interleukin-Resistance (TIR) domain family protein | AT2G03300.1 |
| 2:975590 | 2 | avrRpm | AT2G03030.1 | Toll-Interleukin-Resistance (TIR) domain family protein | | 85070 | 975590 | 27878 | | Toll-Interleukin-Resistance (TIR) domain family protein | AT2G03300.1 |
| 1:11273854 | 1 | avrRpm | | | | | 11273854 | 14598 | RAC1 | Disease resistance protein (TIR-NBS-LRR class) family | AT1G31540.2 |
| 1:11273813 | 1 | avrRpm | | | | | 11273813 | 14639 | RAC1 | Disease resistance protein (TIR-NBS-LRR class) family | AT1G31540.2 |
| 3:2165688 | 3 | avrRpm | | | | | 2165688 | 60264 | RPM1 | NB-ARC domain-containing disease resistance protein | AT3G07040.1 |

**Table S 1:** List of SNPs above the threshold ($-log_{10}(10^{-7})$) and their distance from a given gene set (here, *Arabidopsis thaliana* immune system genes containing NBS-LRR domain)
